# Calcium interaction with Nav1.5 via FGF12A and CaM binding

**DOI:** 10.64898/2026.01.20.700379

**Authors:** Lucy Woodbury, Anna Li, Paweorn Angsutararux, Martina Marras, Emily Wagner, Jonathan R. Silva

## Abstract

Voltage-gated Na^+^ (Nav) channels, including Nav1.5, are responsible for the initiation of cardiac and neuronal action potentials. Regulation of Nav1.5 inactivation is linked to multiple accessory proteins that bind its C-terminal domain (CTD) including calmodulin (CaM) and intracellular fibroblast growth factors (iFGF). Previous results demonstrate that Ca^2+^-bound CaM preferentially binds to iFGF12A. The role of intracellular Ca^2+^ ([Ca^2+^]_i_) in regulating Nav1.5 gating, either directly or via auxiliary proteins like CaM, is controversial. We hypothesize that CaM binding to the Nav1.5 CTD and iFGF12A synergistically alters channel inactivation in a previously unobserved calcium-dependent manner. We performed Fluorescence Resonance Energy Transfer (FRET) imaging in live cells to observe the interaction between the Nav1.5 alpha subunit, CaM and iFGF12A. At resting [Ca^2+^]_i_, a 2-fold difference between acceptor and donor FRET efficiency was observed, implying that a single CaM acceptor is present on the Nav1.5 CTD even in the presence of FGF12A. After increasing [Ca^2+^]_i_, the donor and acceptor FRET efficiencies equalize, suggesting a 2:1:1 ratio between CaM, FGF12A, and the Nav1.5 CTD. We then compared the voltage-dependent gating kinetics of Nav1.5 with FGF12A in the presence/absence of calcium. With low [Ca^2+^]_i_, the steady-state inactivation of Nav1.5 with FGF12A was significantly shifted toward hyperpolarized potential compared to resting [Ca^2+^]_i_. Thus, the FGF12A:CaM complex confers a Ca^2+^-dependent mechanism enabling FGF12A modulates the Nav1.5 steady-state inactivation. Additionally, the ability of multiple subunits to bring CaM to the Nav1.5 CTD implies biological redundancy to prevent major alteration to Nav1.5 inactivation in the absence of CaM.

## Introduction

The voltage-gated sodium (Nav) channel, Nav1.5, initiates the cardiac action potential through the transient inward sodium current (I_Na,T_) and also contributes to the initiation of neuronal action potential. Channel activation occurs within milliseconds, driving the rapid depolarization phase, while subsequent fast inactivation sharply reduces current to prevent excessive prolongation of the action potential. The remaining late Na^+^ current (I_Na,L_) is a key regulator of action potential duration (Han *et al*., 2018). Nav1.5 is encoded by *SCN5A* and comprises four homologous repeats (I–IV), each containing six transmembrane segments (S1–S6). The first four segments constitute the voltage-sensing domain (VSD), with the S4 helix carrying multiple positively charged residues that move in response to membrane depolarization. This conformational change actuates Nav channel gating and regulates sodium influx. Both activation and inactivation are critically coupled to the VSDs of repeats III and IV, and pathogenic variants within these regions that disrupt finely-tuned gating kinetics predispose patients to arrhythmogenic syndromes and epilepsies (Johnson *et al*., 2009; Aurlien *et al*., 2009; Remme & Bezzina, 2010;

Parisi *et al*., 2013; Varga *et al*., 2015; Miyazaki *et al*., 2016; Hsu *et al*., 2017). The III–IV linker, in particular, mediates fast inactivation via the conserved IFM motif, which allosterically modulates the pore domains for fast inactivation (West *et al*., 1992; Jiang *et al*., 2020; Chinthalapudi *et al*., 2024). Cryo-EM structures further demonstrate that the C-terminal domain (CTD) directly interacts with the IFM motif during gating transitions and serves as a scaffold for auxiliary regulatory proteins, including calmodulin (CaM) and intracellular fibroblast growth factors (iFGFs) (Abriel, 2010; Pitt & Lee, 2016; Jiang *et al*., 2020; Biswas *et al*., 2024). CaM in particular is an ubiquitous calcium sensor and has various orientations based on whether it is calcium (Ca^2+^)-saturated (Barbato *et al*., 1992; Zhang *et al*., 2012). Additionally, the A-splice variants of iFGFs contain a noncanonical CaM binding site on their N-terminal (Mahling *et al*., 2021), providing multiple interaction sites for CaM on Nav channel CTDs. These interactions underscore a central question in sodium channel physiology: whether intracellular Ca^2+^ ([Ca^2+^]_i_) modulates Nav1.5 directly, or indirectly through CaM and iFGFs bound to the CTD.

Ca^2+^ is a universal second messenger that plays a central role in both cardiac physiology and neuronal function. In the heart, voltage-gated L-type Ca^2+^ channels initiate excitation–contraction coupling by triggering Ca^2+^ release from the sarcoplasmic reticulum (SR) through ryanodine receptor 2 (RyR2) (Marks, 2013; Sutanto *et al*., 2020). This surge elevates cytoplasmic Ca^2+^ to ∼1 µM, activating muscle contraction and various Ca^2+^-dependent signaling pathways (Berridge, 1998; Sutanto *et al*., 2020). To restore resting concentrations, Ca^2+^ is actively pumped back into the SR by ATPase 2a (SERCA2a), reducing cytosolic concentrations to ∼100 nM (Marks, 2013).

These cyclical fluctuations in intracellular Ca^2+^ are essential for normal cardiac function and continuously reshape the local environment of membrane proteins. Pathogenic changes to calcium cycling within cardiomyocytes, such as chronic atrial fibrillation or sepsis, often lead to delayed afterdepolarizations and arrhythmogenesis (Voigt *et al*., 2012; Hobai *et al*., 2015; Dobrev & Wehrens, 2017; Pinto *et al*., 2017). Beyond the heart, Ca^2+^ also regulates neuronal excitability through a parallel process. [Ca^2+^]_i_ is released from endoplasmic reticulum stores via RyR channels, influencing neurotransmitter release and reuptake with Ca^2+^ waves reaching amplitudes of ∼5 μM (Berridge, 1998; Ross, 2012; Walters & Usachev, 2023).

[Ca^2+^]_i_ concentration is well-known to regulate voltage-gated calcium channels and auxiliary proteins such including CaM, however whether Ca^2+^ directly modulates Nav channels in highly debated. Wingo et al. (2004) first identified an EF-hand–like structure within the Nav1.5 CTD that was hypothesized to directly sense intracellular Ca^2+^ (Anon, n.d.). A subsequent theory suggested that this CTD EF-hand could regulate a Ca^2+^-dependent interaction between calmodulin (CaM) and the downstream IQ motif (∼120 residues away)(Shah *et al*., 2006).

However, later NMR studies provided evidence against this model (Chagot *et al*., 2009). While mutations within the CTD EF-hand region clearly alter Nav1.5 gating (Gardill *et al*., 2018), these effects may not result from direct Ca^2+^ binding. These results shifted attention to indirect Ca^2+^ modulation via auxiliary subunits (Kim *et al*., 2004a). CaM binds the conserved IQ motif on the Nav1.5 CTD and has been implicated in multiple regulatory roles, most notably in suppressing the late sodium current (I_Na,L_) (Deschênes *et al*., 2002; Kim *et al*., 2004a; Yan *et al*., 2017).

Given that CaM is a ubiquitous Ca^2+^ sensor, it was proposed that its Ca^2+^-binding state could influence Nav1.5 gating. Yet, experimental studies have not provided clear evidence for direct Ca^2+^-dependent modulation of Nav1.5 function through CaM binding (Sarhan *et al*., 2012; Ben Johny *et al*., 2013; Ben-Johny *et al*., 2014). Mutagenesis has often been used to probe CaM:CTD interactions, with the IQ(1908–09)/AA substitution serving as a standard approach to disrupt CaM binding. However, this strategy may also alter interactions between the CTD and the DIII–DIV linker IFM motif, complicating interpretation (Johnson, 2020). Importantly, Ben-Johny et al. (2016) demonstrated a fixed 1:1 stoichiometry of CaM bound to Nav1.5, regardless of Ca^2+^ occupancy (Ben-Johny *et al*., 2016). Other studies have suggested that differences in the orientation of apo- versus Ca^2+^-bound CaM may alter how CaM:CTD complexes interact with the DIII–DIV linker (Pitt & Lee, 2016). Thus, a central unresolved question is whether the Ca^2+^-binding state of CaM (apo vs. Ca^2+^-saturated) differentially regulates Nav1.5 gating. Moreover, this raises the possibility that CaM could influence Nav1.5 through non-canonical mechanisms, potentially involving its coordination with other auxiliary subunits at the CTD.

We previously investigated how A-splice variants of intracellular fibroblast growth factors (iFGFs) and CaM regulate voltage-gated sodium channels, focusing on FGF12A interaction with Nav1.5. FGF12A binds the Nav1.5 CTD and is known to suppress the late sodium current (I_Na,L_) (Chakouri *et al*., 2022). All A-splice variants of iFGFs share a conserved CaM-binding domain on their N-terminus (Mahling *et al*., 2021). Our recent work revealed that FGF12A modulates Nav1.5 gating through two distinct mechanisms: (1) a CaM-independent pathway that reduces I_Na,L_, and (2) a CaM-dependent pathway that requires CaM binding to the FGF12A N-terminus to consistently regulate voltage-dependent inactivation (Woodbury *et al*., 2025). CaM must be Ca^2+^-saturated to bind FGF12A (as well as other A-splice variants) (Mahling *et al*., 2021). Thus, Ca^2+^ itself does not directly affect Nav1.5 but rather determines whether CaM can interact with FGF12A. Based on this observation, we hypothesized that changes in [Ca^2+^]_i_ could modulate Nav1.5 gating by dynamically altering the Nav1.5 auxiliary subunit complex makeup via FGF12A and CaM. To test this hypothesis, we designed the present study to determine whether Ca^2+^-dependent changes in Nav1.5:FGF12A:CaM stoichiometry alter Nav1.5 gating. We conducted live-cell FRET imaging and whole-cell manual patch-clamping on transiently transfected HEK293 cells to characterize the Ca^2+^-dependent interactions on Nav1.5 gating.

## Methods

### Molecular Biology and HEK293 culture

All Nav1.5 variants including the IQ1908-1909AA variant and β1-IRES-Nav1.5 variants with and without fluorescent tags were generated in the SCN5A gene using High-Fidelity PCR via the Herculase II Fusion DNA Polymerase (Agilent). All fluorescently tagged CaM plasmids were generated in a similar manner. The various iFGF plasmids and Nav1.5 CTD peptides used were purchased from Vector Builder. Plasmids were amplified from glycerol E. coli stock with NucleoSpin Plasmid DNA purification (Macherey-Nagel).

Magnetic-Assisted Transfection (Magnetofection from Oz Biosciences) was used for all transient transfections of HEK293 cells for FRET and manual patch clamping experiments. For FRET imaging, HEK293 cells were plated at a density of 250k per 35 mm plate 24 hours before transfection. A ratio of 2:1:1 (Nav1.5 channel:FGF12A:CaM) was used to equal a total of 2 μg DNA in 100 μL HBSS buffer with 1 μL Magnetofection reagent per 35 mm plate. For FRET experiments with only two constructs, a ratio of 1:1 was used. Plates were incubated at 37 ° C for 48 hours before imaging occurred.

For whole-cell manual patch-clamping experiments, HEK293 cells were plated at a density of 75k per 35 mm plate 24 hours before transfections. A β1-IRES-Nav1.5 plasmid was used to ensure co-transfection with the auxiliary subunit. 2 μg of the Nav1.5 channel DNA with 1 μg FGF12A DNA was transfected with 200 μL HBSS buffer and 2 μL Magnetofection reagent per 35 mm plate. Plates were incubated at 37 ° C for 48 hours before electrophysiology experiments occurred.

### Electrophysiology and FRET solutions

FRET solutions (in mM): Hank’s balanced salt solution (HBSS, calcium, magnesium, no phenol red): 1.26 CaCl_2_, 0.49 MgCl_2_, 0.41 MgSO_4_, KCl 5.33, 0.44 KH_2_PO_4_, 4.17 NaHCO_3_, 137.93 NaCl, 0.34 NaH_2_PO_4_, 5.56 D-glucose (Thermo Fisher Scientific). Krebs Ringer Buffer (High Ca^2+^): 145 NaCl, 5 KCl, 1.3 MgCl_2_, 1.2 NaH_2_PO_4_, 10 D-glucose, 20 HEPES (pH to 7.4 with NaOH). Extracellular CaCl_2_ concentrations were added at 10 mM. Additionally, 4 μM ionomycin was added immediately before incubation.

Electrophysiology Solutions (in mM): External bath solution: 140 NaCl, 5 KCl, 2 CaCl_2_, 1 MgCl_2_, 10 HEPES (pH to 7.4 with NaOH). Internal pipette solution: 35 NaCl, 105 CsF, 2 MgCl_2_, 10 EGTA, 10 HEPES (pH to 7.2 with CsOH). The final osmolarity of both solutions were between 295-305 mOsm. For low Ca^2+^ measurements, 10 μM BAPTA was added to the internal pipette solution.

### FRET imaging

The FRET experimental protocol was modified from Wang et al. 2023 and Ben-Johnny et. al. 2016. The controls from each experiment included three groups: donor only cerulean-tagged, acceptor only Venus-tagged, and spurious FRET constructs of a Venus-tagged protein and an untagged cerulean. These controls were used to measure experimental variables such as RA1 and RD1 for overall FRET efficiency calculations. Each plate was washed with HBSS(-/-) solution (Thermo Fisher Scientific) twice before being imaged in HBSS solution. For elevated [Ca^2+^]_i_ concentrations, plates were washed and incubated in Krebs Ringer buffer solution with 1 μM ionomycin for 10 minutes at room temperature prior to imaging. The resting Ca^2+^ experiments were to mimic basal levels of [Ca^2+^]_i_ concentrations at diastole. The elevated or “high” Ca^2+^ were to saturate the [Ca^2+^]_i_ levels beyond that of cardiac systole (Kang *et al*., 2023a).

The custom FRET imaging rig is previously described in detail from Wang et. al. 2023. Approximately 50 images for each excitation and emission wavelength paring were collected per plate at a 20X magnification. These pairings included the donor channel: CFP excitation (440 nm) and CFP emission (475 nm), FRET channel: CFP excitation (440 nm) and YFP emission (543 nm), and acceptor channel: YFP excitation (510 nm) and YFP emission (543 nm). All comparisons between measured FRET values were conducted only on groups transfected and imaged on the same day to prevent differences in transfection efficiency or potential shifts in instrument calibration. Using the three measured values, the donor and acceptor centric apparent FRET efficiency values (ED and EA respectively) were calculated with MATLAB using analysis software generate by Wang et al 2023(Kang *et al*., 2023b). The ratio of ED and EA for each cell was calculated to determine the relative ratio of donor to acceptor fluorophores as shown in Ben-Johnny et al. 2016 (Ben-Johny *et al*., 2016). We initially confirmed the relationship between ED/EA to the stoichiometry of auxiliary subunits to full channels using tagged voltage-gated potassium channel KCNQ1 and CaM (**SI Figure 2**). More information is available in the appendix.

**Figure 1:**
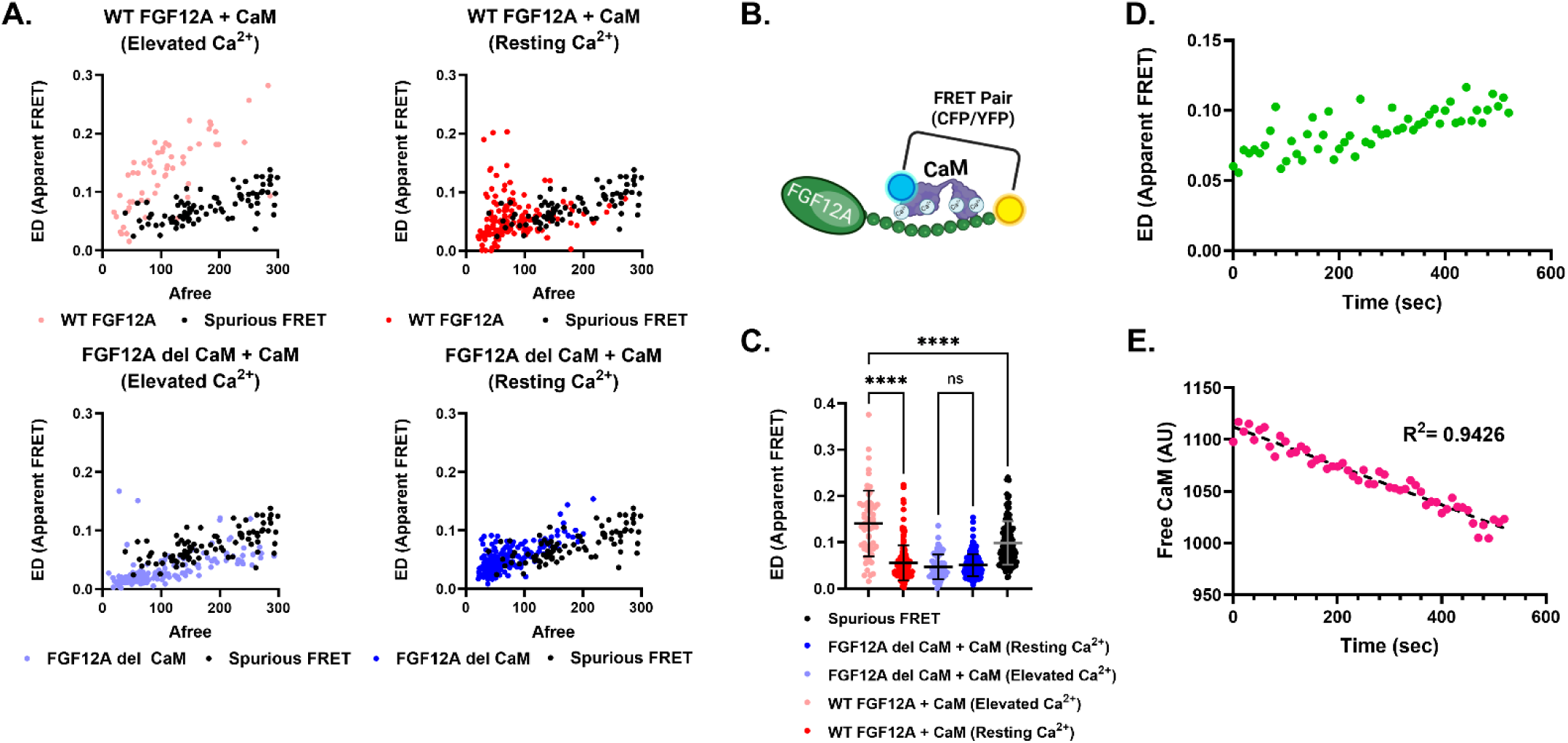
CaM binding to FGF12A is calcium-dependent. **(A)** In elevated [Ca^2+^]j, WT FGF12A shows strong interaction to CaM *{red),* while the CaM-binding deficient mutant *{FGF12A del CaM, blue)* shows no significant interaction compared to background FRET *{black).* (B) Schematic of the FRET donor-acceptor pair. (C) Apparent FRET efficiency (ED) significantly increases for WT FGF12A in elevated versus resting [Ca^2+^]j (**‘*p < 0.0001), but not for FGF12A del CaM. (D) Free CaM (unbound acceptor) decreases over time after [Ca^2+^]j elevation. (E) Correspondingly, ED increases over time as CaM becomes bound to FGF12A.

**Figure 2:**
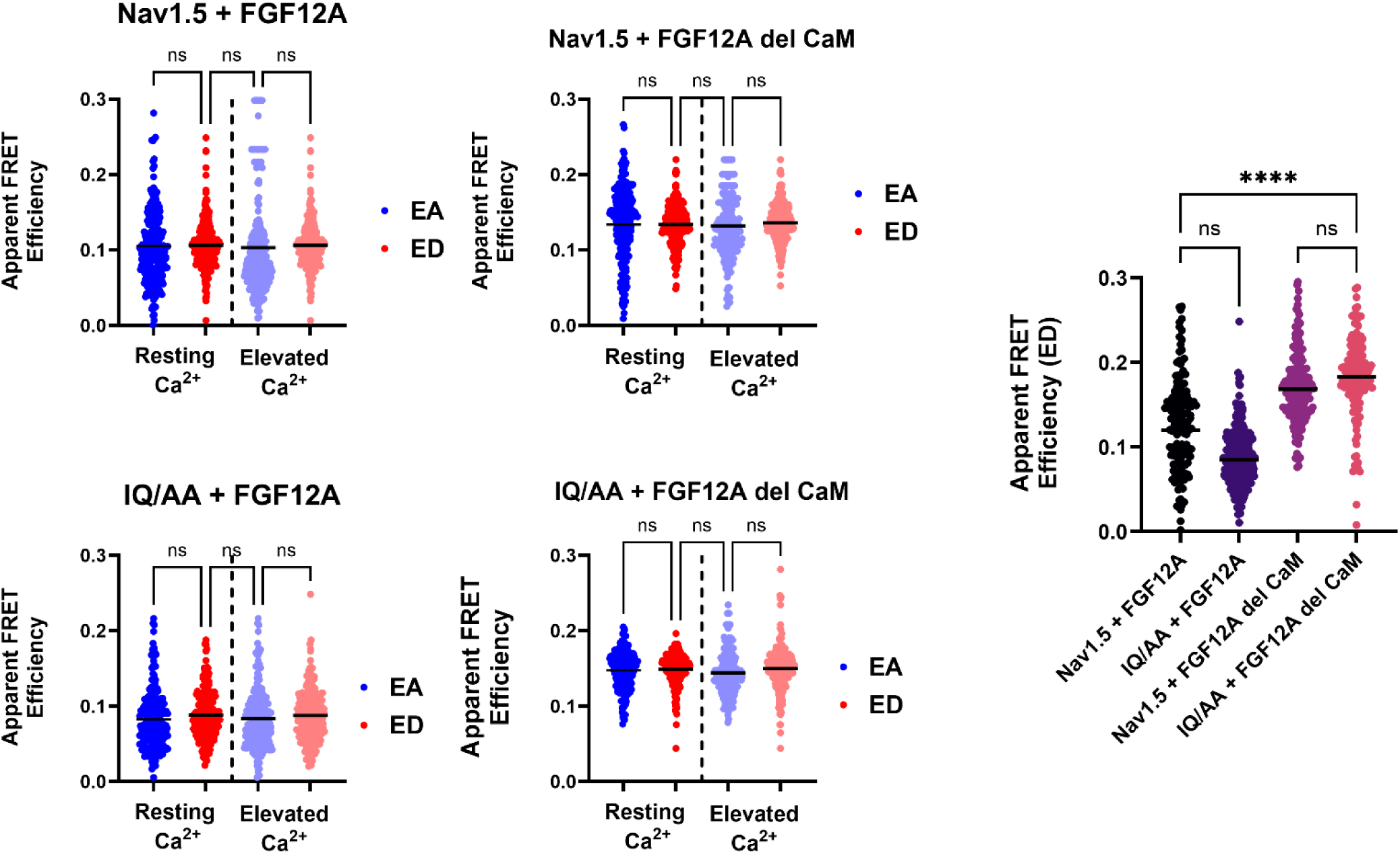
Ratio analysis of FGF12A-Nav1.5 interactions. Scatter dot plots (mean shown) reveal no significant differences between donor- (ED) and acceptor-centric (EA) FRET efficiencies across conditions, were consistent with a 1:1 ratio of Navi .5 to FGF12A regardless of CaM binding or [Ca^2+^]j levels. The summary plot (right) highlights a significant increase in ED for FGF12A del CaM compared to WT FGF12A when bound to Navi .5 (****p < 0.0001).

### Whole-Cell Manual Patch Recordings

The initial protocol steps were developed from Angsutararux et al. 2023 (Angsutararux *et al*., 2023). Whole-cell manual patch-clamp recordings were collected from transiently transfected HEK293 cells no older than passage 20. A custom patch-clamp rig was used which consisted of an Axopatch 200B (Molecular Devices) amplifier with a Digidata 1440A (Molecular Devices) acquisition system on the pClamp 10 (Molecular Devices) software. Pipette resistances were constrained between 2-4 MΩ when filled with the internal pipette solution. Individual cells were chosen, and a gigaohm-seal (> 1 GΩ) was used to confirm the whole-cell clamp configuration. Cells were allowed 5 minutes to establish equilibrium between the cytosol and internal pipette solution. Using a holding potential of -70 mV, the whole-cell membrane capacitances (C_m_), input resistances (R_in_), and series resistance (Rs) were measured and electronically compensated. For elevated Ca^2+^ measurements, plates were incubated at room temperature in the Krebs-Ringer Buffer (High Ca^2+^) solution + 4 μM ionomycin for 10 minutes prior to recording with control external (bath) and internal (pipette) solutions. For low Ca^2+^ measurements, plates were incubated at 37 ° C for three hours in calcium-free HEK cell media, prior to recording with control external (bath) solution and reduced Ca^2+^ internal (pipette) solution + 10 μM BAPTA. Cells were held at a holding potential of -70mV for 10 minutes after cell membrane rupture to ensure equilibrium of calcium levels. Electrophysiology data was recorded in Clampfit (Molecular Devices) and analyzed using Jupyter Notebook and Prism 10 (GraphPad). P values were evaluated using one-way Welch ANOVA unpaired t-test or Tukey’s multiple comparisons test.

## Results

### FGF12A binding to CaM depends on [Ca^2+^]_i_ concentration

To recapitulate previous reports that showed FGF12A association with CaM using purified proteins (Mahling *et al*., 2021), we measured the apparent FRET efficiency between WT FGF12A and CaM under resting and elevated intracellular Ca²⁺ conditions with our live-cell FRET imaging system (**Figure 1**). CaM was tagged with Cerulean (CFP, donor) and FGF12A with Venus (YFP, acceptor), fluorophore assignments that remain consistent throughout unless otherwise noted (**Figure 1B**). Throughout these experiments, the donor-centric (ED) and acceptor-centric (EA) values were equivalent (**Figure A3**), therefore we are only reporting ED here for consistently and clarity, both values can be found in **Table 1**. At elevated Ca²⁺, ED increased significantly compared to resting Ca²⁺ (+0.085 AU, ±0.006, p < 0.0001), in agreement with work by Mahling et al. (2021), which reported that FGF12A binds only Ca²⁺-saturated CaM (**Figures 2A, 2C**). Importantly, this increase also exceeded spurious FRET levels (0.042 AU, ±0.006, p < 0.0001; see Methods), implying that the FRET measurement reflected genuine FGF12A:CaM interaction rather than random collisions. As the ionomycin treatment to increase the [Ca^2+^]_i_ concentration as described in the methods requires a 10-minute incubation, we performed time-lapse imaging of WT FGF12A:CaM interactions during this process. As [Ca^2+^]_i_ levels rose, ED values increased steadily (**Figure 1E**), while the pool of free CaM (AU) declined in parallel (**Figure 1D**). This inverse relationship indicates that CaM progressively binds FGF12A as intracellular Ca²⁺ increases.

**Table 1:** Resting and Elevated Calcium EA and ED of CaM binding partners.

|  | *EA (Resting Ca<sup>2+</sup>)* | *ED (Resting Ca<sup>2+</sup>)* | *EA (Elevated Ca<sup>2+</sup>)* | *ED (Elevated Ca<sup>2+</sup>)* | *N* |
| --- | --- | --- | --- | --- | --- |
| *FGF12A-YFP + CaM-CFP* | 0.064 ± 0.006 | 0.074 ± 0.005 | 0.141 ± 0.011 | 0.1404 ± 0.009 | 162 |
| *FGF12A del CaM-YFP + CaM-CFP* | 0.039 ± 0.002 | 0.042 ± 0.002 | 0.039 ± 0.004 | 0.047 ± 0.003 | 207 |
| *Nav1.5-YFP + FGF12A-CFP* | 0.0951 ± 0.005 | 0.107 ± 0.008 | 0.103 ± 0.003 | 0.105 ± 0.002 | 164 |
| *Nav1.5-YFP + FGF12A del CaM-CFP* | 0.134 ± 0.003 | 0.133 ± 0.002 | 0.132 ± 0.003 | 0.134 ± 0.002 | 171 |
| *IQ/AA-YFP + FGF12A-CFP* | 0.083 ± 0.003 | 0.087 ± 0.002 | 0.077 ± 0.003 | 0.087 ± 0.003 | 254 |
| *IQ/AA-YFP + FGF12A del CaM-CFP* | 0.148 ± 0.002 | 0.149 ± 0.002 | 0.114 ± 0.003 | 0.149 ± 0.002 | 144 |
| *Nav1.5-YFP + CaM-CFP* | 0.129 ± 0.011 | 0.130 ± 0.003 | 0.139 ± 0.012 | 0.127 ± 0.006 | 200 |
| *IQ/AA-YFP + CaM-CFP* | 0.062 ± 0.009 | 0.067 ± 0.003 | 0.052 ± 0.006 | 0.057 ± 0.004 | 107 |

To disrupt CaM binding, we used a mutant construct lacking the canonical CaM-binding region (residues 33–61; FGF12A del CaM) (Woodbury *et al*., 2025). In this case, no measurable interaction was detected at any Ca²⁺ concentration above spurious FRET (**Figures 1A, 1C**). While Mahling et al. (2021) suggested that the N-terminal long-term inactivation particle (residues 1–30) might weakly interact with Ca²⁺-bound CaM, we observed no such binding in our system. Instead, at elevated Ca²⁺, WT FGF12A displayed a strict 1:1 stoichiometry with CaM, supported by equivalent EA and ED values (**Figure A3B**). Thus, our results demonstrate that in a heterologous expression system, WT FGF12A binds a single Ca²⁺-saturated CaM molecule exclusively at its designated alpha subunit CaM-binding site.

### FGF12A binds to Nav1.5 with 1:1 stoichiometry

Having established the interaction between CaM and FGF12A, we next examined whether CaM influences FGF12A binding to Nav1.5. Intracellular FGFs (iFGFs) are known to bind the Nav1.5 C-terminal domain (CTD) at residues E1890–R1898(Wang *et al*., 2011, 2012) in close proximity to the CaM-binding site at residues 1908–1909 (Bähler & Rhoads, 2002; Tan *et al*., 2002; Pitt & Lee, 2016). To test whether CaM affects this interaction, we performed FRET imaging in transiently transfected HEK cells, using a 2:1 DNA plasmid ratio of Nav1.5 to FGF12A to ensure robust channel expression. As shown in **Figure 2**, donor- and acceptor-centric FRET efficiencies (ED and EA) were equivalent under both resting and elevated Ca²⁺ conditions. This pattern was consistent across all tested combinations: WT Nav1.5 with WT FGF12A, WT Nav1.5 with FGF12A del CaM, IQ/AA Nav1.5 with WT FGF12A, and IQ/AA Nav1.5 with FGF12A del CaM (**Table 1**). Therefore, regardless of Ca²⁺ concentration or the ability of Nav1.5 or FGF12A to bind CaM, the stoichiometry of the Nav1.5:FGF12A interaction remains fixed at 1:1.

Unexpectedly, FGF12A del CaM showed significantly high apparent FRET efficiency (ED) with Nav1.5 compared to WT FGF12A (**Figure 2, right**). The FGF12A del CaM mutant produced higher FRET efficiencies with both WT Nav1.5 (+0.065 AU ± 0.009, p < 0.0001) and IQ/AA Nav1.5 (+0.076 AU ± 0.008, p < 0.0001). This increased binding may reflect structural changes in FGF12A caused by removal of its CaM-binding domain, potentially altering its conformation or accessibility to Nav1.5.

### CaM interacts with Nav1.5 comparably to FGF12A

Figure 3 summarizes the interactions of CaM with all binding partners under resting and elevated Ca²⁺ conditions. Consistent with Figure 1, WT FGF12A showed a significant increase in donor-centric FRET efficiency (ED) when Ca²⁺ was elevated, whereas FGF12A del CaM displayed no change (**Table 1**). To test whether this Ca²⁺ dependence was unique to FGF12A, we compared CaM binding to Nav1.5. Unlike FGF12A, WT Nav1.5 exhibited no significant difference in ED between resting and elevated Ca²⁺, despite known conformational differences between apo- and Ca²⁺-saturated CaM (Barbato *et al*., 1992; Pitt & Lee, 2016). As expected, the IQ(1908–09)/AA mutation abolished CaM binding to the Nav1.5 CTD under all conditions (Figure 3).

**Figure 3:**
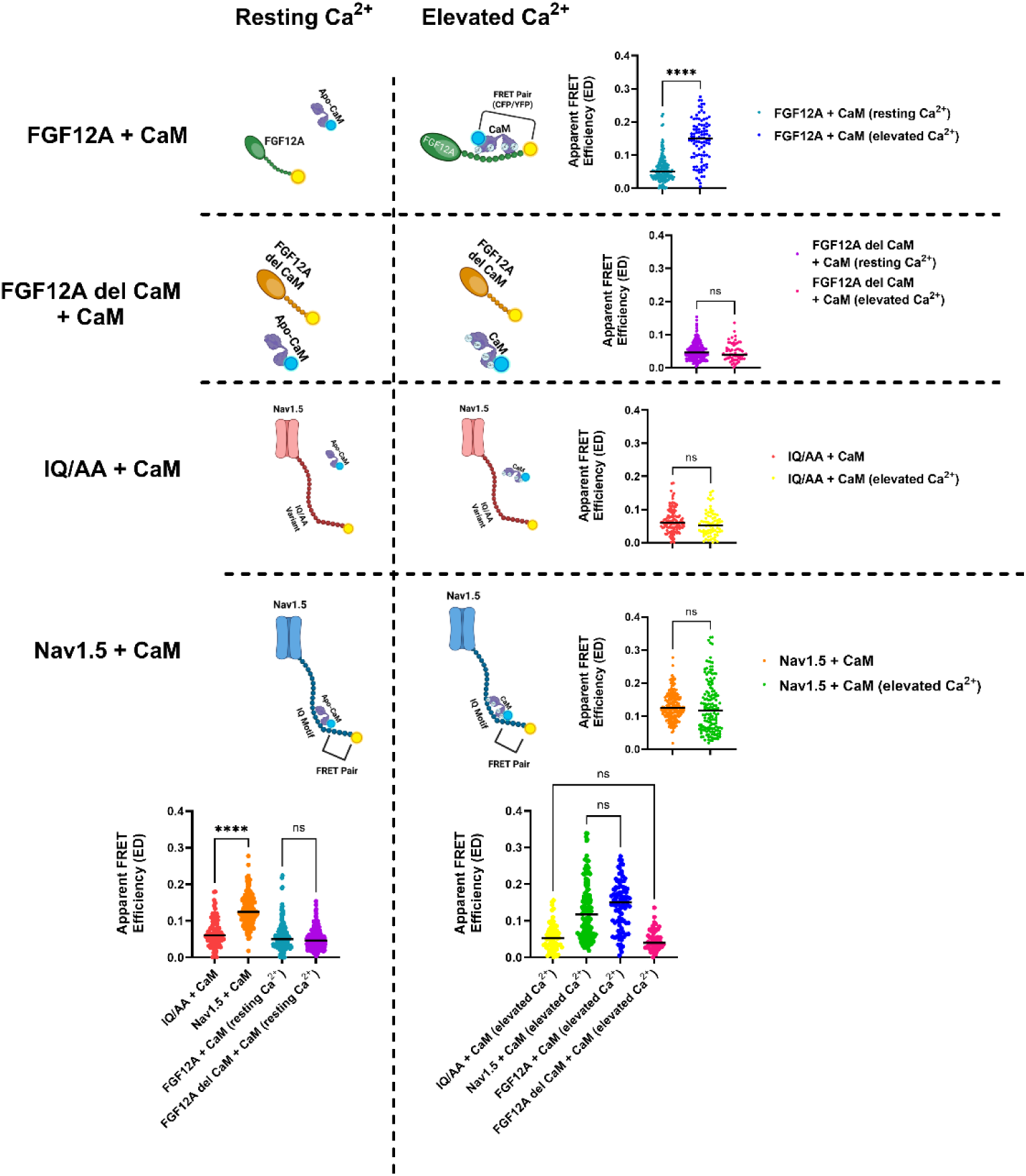
Comparative analysis of CaM binding partners across calcium conditions. Each row shows how individual protein pairs respond to changing [Ca^2+^]j, while each column compares different pairs under the same condition. **Rows:** Only WT FGF12A + CaM display a calcium-dependent increase in donor-centric FRET efficiency (ED) (****p < O.ŨŨO1). No significant Ca^2+^-dependent changes were observed for FGF12A del CaM + CaM, IQ/AA + CaM, or WT Navi .5 + CaM. **Columns:** At resting Ca^2+^, ED values are similarly low for WT FGF12A + CaM, FGF12A del CaM + CaM, and IQ/AA + CaM, while WT Navi .5 + CaM shows a significantly higher ED than IQ/AA Nav1.5 + CaM (***‘p < 0.0001). At elevated Ca^2+^, ED values for WT FGF12A + CaM and WT Navi .5 + CaM are comparable to their respective controls. Overall, significant differences were observed only between resting vs. elevated Ca^2+^ for WT FGF12A + CaM, and between WT and IQ/AA Navi .5 + CaM at resting Ca^2+^.

When comparing protein configurations at resting Ca²⁺, only WT Nav1.5:CaM interactions showed an increase in ED (+0.051 AU ± 0.005, p < 0.0001) relative to the IQ/AA mutant. At this baseline, no significant difference was observed between WT FGF12A and FGF12A del CaM interactions with CaM (Figure 3). Under elevated Ca^2+^, only WT FGF12A:CaM and WT Nav1.5:CaM maintained higher ED values compared to their inhibited counterparts (FGF12A del CaM and IQ/AA Nav1.5). Notably, there was no significant difference in ED between WT FGF12A:CaM and WT Nav1.5:CaM, indicating that Ca-^2+^saturated CaM does not preferentially bind one partner over the other. Together, these results suggest that at elevated [Ca^2+^]_i_, CaM can independently and equivalently associate with either FGF12A or the Nav1.5 CTD.

### [Ca^2+^]_i_ increases enable multiple CaM proteins to bind to the Nav1.5:FGF12A complex

In **Figure A2** and the appendix, we demonstrated that the ratio of donor- and acceptor-centric FRET efficiencies (ED/EA) reflects the stoichiometric ratio of donor to acceptor fluorophores. We next applied this approach to quantify the number of CaM molecules bound to the Nav1.5 CTD in the presence of WT FGF12A at different Ca^2+^ concentrations. Because both Nav1.5:CaM and FGF12A:CaM interactions were equivalent at elevated Ca^2+^ (Figure 3), we designated the two YFP-tagged proteins (Nav1.5-YFP and FGF12A-YFP) as acceptors and CaM-CFP as the donor. At resting [Ca^2+^]_i_, ED/EA values indicated a 2:1 acceptor-to-donor ratio (Figure 4**-1**), consistent with a single CaM molecule bound to the Nav1.5 CTD, likely in its apo form. When [Ca^2+^]_i_ was elevated, this ratio shifted to 2:2 (1:1) (Figure 4**-2**), suggesting that two CaM molecules were now positioned within the Nav1.5:FGF12A complex. To determine which binding partner recruits the second CaM, we repeated the experiments with all combinations of WT FGF12A, FGF12A del CaM, WT Nav1.5, and IQ/AA Nav1.5. Figure 4 summarizes these results, with each labeled corner representing a specific Nav1.5:FGF12A pair under resting or elevated Ca^2+^ conditions. Color-coded arrows highlight key comparisons: *black* (WT vs. del CaM FGF12A), *red* (WT vs. IQ/AA Nav1.5), and *blue* (resting vs. elevated Ca^2+^).

**Figure 4:**
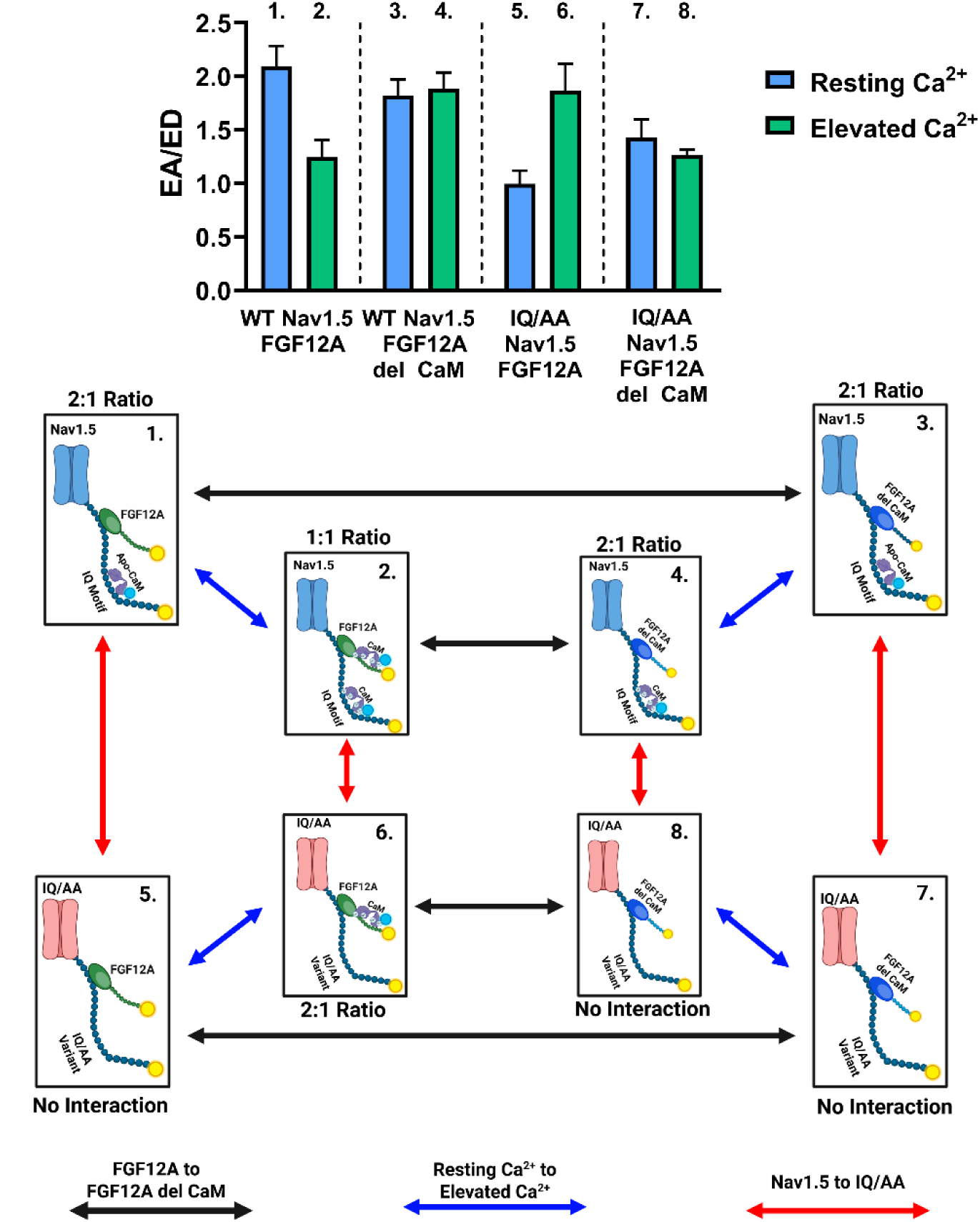
Model of auxiliary protein complexes illustrating stoichiometric changes between labeled donors and acceptors. Complexes were formed using combinations of WT or IQ/AA Nav1.5, WT FGF12A or FGF12A del CaM, and WT CaM, and imaged under resting or elevated Ca^2+^. Black arrows indicate substitutions of FGF12A, red arrows substitutions of Nav1.5, and blue arrows changes in Ca^2+^ conditions. The donor-acceptor ratios (CaM-CFP donors vs. Nav1.5-YFP and FGF12A-YFP acceptors) were experimentally determined, with each label directly linked to its corresponding graph segment.

Replacing WT FGF12A with FGF12A del CaM abolished the Ca^2+^-dependent shift in donor proteins. At resting Ca^2+^, the ED/EA ratio remained 2:1 (**Figure 4-3**), consistent with a single CaM bound to Nav1.5:FGF12A del CaM. Unlike with WT FGF12A, however, raising the Ca^2+^ concentration did not recruit an additional CaM (Figure 4**-4**), indicative that the second CaM molecule normally associates with WT FGF12A only. Conversely, when CaM binding to the Nav1.5 CTD was inhibited with the IQ/AA mutation, no significant interaction was detected at resting Ca^2+^ (**Figure 4-5, SI Figure 4, Table 2**), as expected. Upon [Ca^2+^]_i_ elevation with the IQ/AA variant, the ED/EA ratio shifted to 2:1 from no measurable interaction (**Figure 4-6**), reflecting two acceptors (IQ/AA Nav1.5 and WT FGF12A) bound to a single CaM. Since IQ/AA Nav1.5 cannot bind CaM beyond random fluctuations (Figure 3), this interaction must originate from CaM binding to WT FGF12A. Finally, when both binding sites were disrupted (IQ/AA Nav1.5 + FGF12A del CaM), no measurable interaction was observed at any Ca^2+^ concentration (Figure 4**-7**, Figure 4**-8****, SI** Figure 4, **Table 2**). Together, these findings indicate that at resting Ca^2+^, CaM primarily binds the Nav1.5 CTD, while at elevated Ca^2+^, a second CaM molecule engages WT FGF12A, establishing a dual binding configuration within the Nav1.5:FGF12A complex.

**Figure 5:**
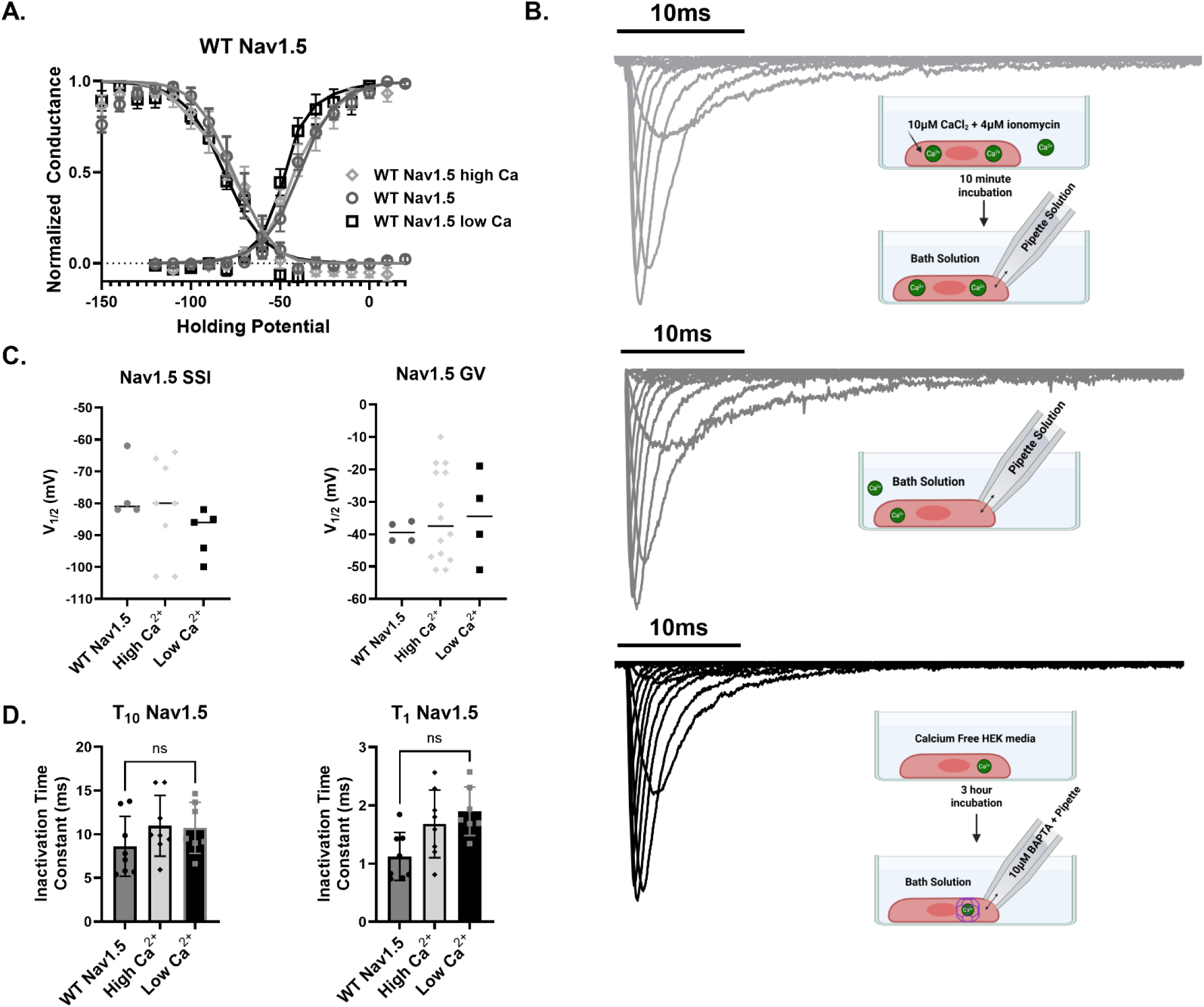
WT Nav1.5 function is unaffected by [Ca^2+^]j. All error bars represent SEM. (A) GV and SSI curves for WT Nav1.5 expressed without FGF12A or FGF12A del CaM *{diamonds: high [Ca^2+^]i, circles: control, squares: low [Ca^2+^]_j_’)* show no calcium-dependent shifts. (B) Representative whole-cell patch-clamp traces with protocol schematic *{light gray: high [Ca^2+^]j, gray: control, black: low [Ca^2+^]j).* **(C)** Scatter plots of SSI V_1/2_ (left) and GV V_1/2_ (right) with mean values indicate no significant differences across calcium conditions. **(D)** Inactivation time constants (τ_10_ and Tļ) show no significant changes with altered [Ca^2+^]j.

**Figure 6:**
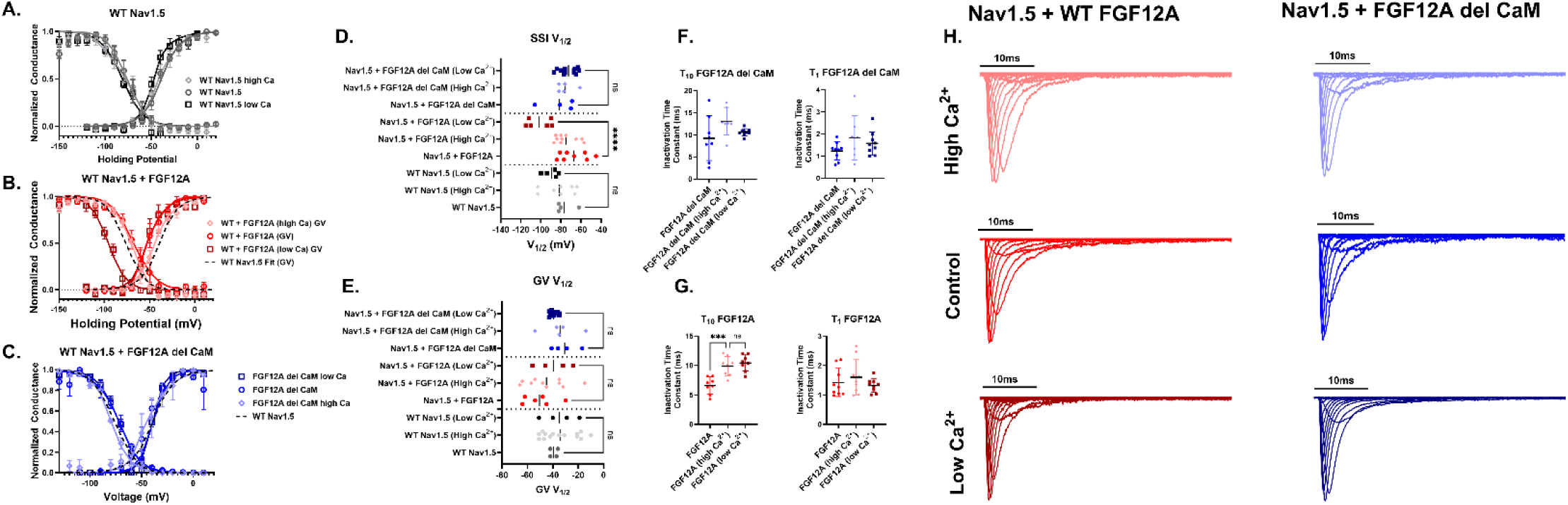
Calcium-dependent effects of FGF12A on Nav1.5 gating. HEK293 cells were transiently transfected with Nav1.5, β1, and either WT FGF12A or FGF12A del CaM and recorded using manual whole-cell patch clamp at high [Ca^2+^]j *{diamonds’),* control *(circles),* or low [Ca^2+^]j (squares). All error bars are ±SEM. (A) GV and SSI curves of WT Nav1.5 alone (data repeated from Fig. 5A) show no calcium-dependent shifts. (B) With WT FGF12A, SSI is depolarized at control and high [Ca^2+^]j but significantly hyperpolarized at low [Ca^2+^]i. (C) With FGF12A del CaM, no significant GV or SSI shifts were detected at any calcium level. (D) Scatter plot of SSI V^2 shows that only WT FGF12A under low [Ca^2÷^]i produces a significant shift (****p < 0.0001). (E) Scatter plot of GV V_1/2_ reveals no significant calcium-dependent differences. (F) Inactivation time constants (τ_10_ and η) for FGF12A del CaM show no changes. (G) With WT FGF12A, lowering calcium significantly prolongs τι_0_ compared to control (***p = 0.0006). (H) Representative normalized sodium current traces.

**Table 2:** Comparison between FGF12A and FGF12A del CaM FRET efficiency to Nav1.5.

|  | *EA (Resting Ca<sup>2+</sup>)* | *ED (Resting Ca<sup>2+</sup>)* | *EA (Elevated Ca<sup>2+</sup>)* | *ED (Elevated Ca<sup>2+</sup>)* | *N* |
| --- | --- | --- | --- | --- | --- |
| *Nav1.5 + FGF12A* | 0.081 ± 0.006 | 0.129 ± 0.008 | 0.108 ± 0.006 | 0.106 ± 0.007 | 57 |
| *Nav1.5 + FGF12A del CaM* | 0.079 ± 0.004 | 0.111 ± 0.006 | 0.071 ± 0.004 | 0.103 ± 0.005 | 89 |
| *IQ/AA + FGF12A* | 0.043 ± 0.002 | 0.041 ± 0.002 | 0.089 ± 0.004 | 0.116 ± 0.006 | 95 |
| *IQ/AA + FGF12A del CaM* | 0.043 ± 0.002 | 0.041 ± 0.002 | 0.044 ± 0.002 | 0.067 ± 0.005 | 99 |

### WT Nav1.5 is not independently affected by changes in [Ca^2+^]_i_

Having observed a Ca^2+^-dependent shift in the Nav1.5 auxiliary subunit complex, we next asked whether this structural change translated into altered channel gating. Consistent with prior studies (Ben-Johny *et al*., 2014; Niu *et al*., n.d.) varying intracellular Ca²⁺ levels did not measurably affect Nav1.5 gating kinetics (Figure 5) when it was expressed alone. Whole-cell patch-clamp recordings were performed in transiently transfected HEK293 cells under three conditions: elevated Ca²⁺ (*light grey*), control (*dark grey*), and low Ca²⁺ (*black*) (Figure 5B). Elevated Ca²⁺ was achieved by incubating cells for 10 minutes in Krebs-Ringer solution supplemented with 10 μM CaCl₂ and 4 μM ionomycin, matching the FRET protocol. Low Ca²⁺ was induced by incubating cells for three hours in calcium-free media. To buffer intracellular Ca²⁺, the pipette solution contained 10 μM BAPTA (Deschênes *et al*., 2002). Across conditions, neither steady-state inactivation (SSI) nor conductance-voltage (GV) relationships were altered (Figures 5A**, 5C**). Likewise, inactivation kinetics remained unchanged, with no detectable differences in decay time constants at either 10 ms (τ₁₀) or 1 ms (τ₁) (Figure 5D).

### The addition of FGF12A dictates a Ca^2+^-dependent response in the voltage dependence of inactivation

As the Nav1.5 channel independently did not have a Ca^2+^ dependent change in gating kinetics, we continued onto determining whether the changes in the auxiliary subunit complex would alter Nav1.5 gating. We repeated the patch-clamp experiments (Figure 5) with co-transfection of either WT FGF12A (*red*) or FGF12A del CaM (*blue*) alongside Nav1.5 (Figure 6H). As before, recordings were performed under three Ca²⁺ conditions: elevated (*light red/light blue*), control (*red/blue*), and low (*dark red/dark blue*). For cells expressing FGF12A del CaM, no changes were observed in SSI, GV, or inactivation kinetics across conditions (Figures 6C**–F**), consistent with our FRET results showing loss of Ca²⁺ dependence. In contrast, WT FGF12A produced clear Ca²⁺-dependent effects. At low Ca²⁺, Nav1.5 exhibited a significant hyperpolarizing shift in SSI relative to control conditions (–30.25 mV ± 7.72, p = 0.0001) (Figures 6B**, 6D**). Additionally, altering Ca²⁺ levels in the presence of WT FGF12A increased the τ₁₀ inactivation time constant (3.31 ms ± 0.74, p = 0.0006). These results indicate that Ca²⁺-induced changes in Nav1.5 gating kinetics require WT FGF12A and are regulated through the formation of the FGF12A:CaM complex.

## Discussion

We identified a Ca^2+^-dependent mechanism by which FGF12A and CaM modulate Nav1.5 channel gating. Live-cell FRET imaging supported a Ca²⁺-dependent interaction between FGF12A and CaM. Subsequent experiments demonstrated that CaM is associated with both FGF12A and Nav1.5. To quantify this interaction, we analyzed the ratio of donor- and acceptor-centered apparent FRET efficiencies (ED and EA, respectively), which allowed us to assess the stoichiometry of CaM bound to the Nav1.5:FGF12A complex under resting and elevated Ca²⁺ conditions. These analyses revealed an increased number of CaM proteins associated with the Nav1.5 complex at higher Ca²⁺ concentrations. Given this Ca²⁺-dependent structural modulation, we next examined its functional consequence on channel activity. Whole-cell patch-clamp recordings demonstrated that, in the presence of FGF12A, Nav1.5 inactivation becomes sensitive to intracellular Ca²⁺ levels via a CaM interaction with FGF12A.

### Previous Calcium Regulation of Nav1.5

Historically, reports on the modulation of the Nav1.5 channel by [Ca^2+^]_i_ have been conflicting. Initial studies focused on the E-F hand present on the Nav1.5 CTD, but both structural and functional analyses suggested that [Ca^2+^]_i_ had no effect on Nav channels (Chagot *et al*., 2009). Our findings, which examined the direct interaction of [Ca^2+^]_i_ with Nav1.5, confirmed that Nav1.5 independently is likely not directly sensitive to changes in [Ca^2+^]_i_ concentration (Figure 5). When considering auxiliary subunit interactions, CaM was able to bind to the Nav1.5 IQ motif at a 1:1 ratio, regardless of varying [Ca^2+^]_i_ concentrations (Ben-Johny *et al*., 2016). Some researchers found that changes in [Ca^2+^]_i_ levels and alterations to the IQ motif affected stable inactivation and the percentage of late sodium current, indicating a role for CaM (Kim *et al*., 2004b; Yoder *et al*., 2019). However, other studies showed no direct regulation of CaM by [Ca^2+^]_i_ (Ben-Johny *et al*., 2014). Our results did not reveal any significant difference in the interaction between the Nav1.5 CTD and CaM at either resting or elevated [Ca^2+^]_i_ levels (**Figure A3**). Major differences in Nav gating associated with the CaM protein typically depend on its binding to the Nav CTD irrespective of [Ca^2+^]_i_. This interaction can be altered through mutations in the IQ motif or CaM, such as G114R or G114W (Brohus *et al*., 2023). CaM’s presence on the Nav CTD is thought to regulate the kinetics of the DIII-DIV linker, promoting fast inactivation of the channel and inhibiting pathogenic late sodium current (Bähler & Rhoads, 2002; Tan *et al*., 2002; Deschênes *et al*., 2002; Herzog *et al*., 2003; Gabelli *et al*., 2014; Yan *et al*., 2017; Gade *et al*., 2019).

This result raises the intriguing question of whether other auxiliary subunits interacting with CaM in a Ca^2+^-dependent manner might regulate Nav1.5 channel gating. Our previous work indicated that A-splice variants of iFGFs employ two mechanisms to alter Nav channel gating (Woodbury *et al*., 2025). Canonically, A-splice variants are thought to inhibit pathogenic late sodium current (Dover *et al*., 2010; Chakouri *et al*., 2022; Woodbury *et al*., 2025). We observed a CaM-independent mechanism, where CaM does not need to bind to FGF12A, nor does FGF12A require the CaM-binding region, to reduce pathogenic late sodium current. However, we also found that FGF12A requires a bound CaM on its N-terminus to alter the voltage-dependence of Nav1.5 inactivation (Woodbury *et al*., 2025). Moreover, FGF12A requires CaM to be Ca^2+^-bound to interact with its N-terminus (Figure 1). We continued to investigate how this Ca^2+^-dependent interaction might regulate Nav1.5 inactivation. Our study revealed two key findings: (*1*) changes in [Ca^2+^]_i_ concentration can alter the number of CaM proteins within the Nav1.5:FGF12A complex and (*2*) the presence of FGF12A on the Nav1.5 CTD drives Ca^2+^-dependent modulation of Nav1.5 inactivation.

### Changes to the Sodium Channel Complex

We proposed the model shown in Figure 4 using assumptions and results from previous figures and supplementary information. Our initial work confirmed the interaction between Ca^2+^-saturated CaM and FGF12A, as previously detailed by Mahling et al., 2021 (Mahling *et al*., 2021) (Figure 1). We then tested whether the Ca^2+^- dependent interaction between FGF12A and CaM would function within the Nav1.5 CTD complex, or if competition between CaM and other binding partners (*FGF12A and Nav1.5*) would interfere. To evaluate this hypothesis, we examined each aspect of the three-way relationship. We confirmed that each pair of binding partners (*FGF12A:CaM, CaM:Nav1.5, Nav1.5:FGF12A)* interacted in a 1:1 ratio (Figure 2 **and Figure A3**). Next, we assessed whether the presence of CaM on the Nav1.5 CTD affected FGF12A binding to Nav1.5 (Figure 2). Finally, we determined if we could assign CaM location to the two putative binding sites: FGF12A and the Nav1.5 CTD. As shown in Figure 3, under similar experimental conditions, fluorescently tagged FGF12A and CaM exhibited equivalent apparent FRET efficiencies compared to tagged CaM and Nav1.5. The reported dissociation constant (K_d_) values of CaM with Nav1.5 are approximately 105 ± 15 nM, within the range of the K_d_ value of 107 nM for Ca^2+^-saturated CaM with FGF12A (Gabelli *et al*., 2014; Mahling *et al*., 2021). Interestingly, other A-splice variants of iFGFs exhibit a higher affinity (lower K_d_ values) for Ca^2+^-saturated CaM (Mahling *et al*., 2021). This difference in CaM binding affinity between iFGFs suggests that CaM’s binding preference between the Nav1.5 CTD and the N-terminus of other A-splice iFGFs could shift depending on whether CaM is in its apo or Ca^2+^-bound form. Examples of this include FGF14A, which interacts with Nav1.1 in neurons and Nav1.5 in adrenal chromaffin cells (Lou *et al*., 2005; Martinez-Espinosa *et al*., 2021).

Thus, under WT conditions, high [Ca^2+^]_i_ will increase the number of CaM proteins present on the Nav1.5:FGF12A complex. Accordingly, two CaM proteins can simultaneously bind to the Nav1.5 CTD: one to the Nav1.5 IQ motif and another to the N-terminus of FGF12A. However, we cannot determine the specific orientations of the CaM proteins bound to the Nav1.5 complex as CaM can bind in various orientations, including with either lobe or both lobes tightly bound (Zhang *et al*., 2012). Furthermore, we cannot ascertain whether the initial CaM on the Nav1.5 CTD changes or shifts from apo-CaM to Ca^2+^-saturated CaM. Therefore, our research focused on testing whether the Ca^2+^-dependent presence of CaM on the FGF12A N-terminus alters Nav1.5 inactivation, rather than investigating any structural changes in CaM itself.

### Functional Results of Calcium Regulation

Our results reveal a significant change in the voltage dependence of Nav1.5 inactivation when FGF12A is present, specifically at low [Ca^2+^]_i_ concentrations. We observed a dramatic hyperpolarizing shift in steady-state inactivation (SSI) at low [Ca^2+^]_i_ levels with WT FGF12A (Figure 6B). This Ca^2+^ sensitivity is not evident in other configurations, such as Nav1.5 alone or Nav1.5 with FGF12A del CaM. In Figure 6, a hyperpolarizing shift in steady-state inactivation of the Nav1.5:FGF12A complex was noted when [Ca^2+^]_i_ was reduced. Under these conditions, we assume no CaM proteins are bound to FGF12A, though they remain on the Nav1.5 complex based on our stoichiometric calculations (Figure 4). This scenario mimics the behavior of Nav1.5 with the IQ/AA mutation, where no CaM proteins are present with its drastic SSI hyperpolarizing shift (Kang *et al*., 2021; Woodbury *et al*., 2025). This hyperpolarizing shift in steady-state inactivation is absent in other cases where CaM isn’t bound to the FGF12A N-terminus, such as in experiments with FGF12A del CaM or with other iFGFs like FGF12B. Angsutararux et al. suggested that the core domain of iFGFs contributes to differences in voltage dependence of inactivation, as seen between FGF12B and FGF13VY(Angsutararux *et al*., 2023). However, since FGF12A and FGF12B share the same core region (Biadun *et al*., 2024), the core region alone doesn’t account for these differences. The Ca^2+^-dependent shift in steady-state inactivation likely results from the combined effects of the core region and the N-terminus. Mutations within the FGF12A N-terminus, such as V52H, can have significant downstream effects on ion channel function (Huang *et al*., 2024). Therefore, this functional Ca^2+^ dependence on steady-state inactivation must be attributed to the CaM-binding region of FGF12A itself.

Given the observed Ca^2+^-dependent effect, it is pertinent to question why Nav1.5 channel inactivation remains unaffected at resting or basal levels of [Ca^2+^]_i_ when co-transfected with WT FGF12A. This is especially intriguing as no significant interactions between CaM and FGF12A were observed under similar experimental conditions (Figure 1). One potential explanation could be the limitations in the precision of our FRET measurements, which may not accurately capture all interactions between the proteins. Our methodology does not allow for the direct calculation of the ratio of Ca^2+^-saturated CaM to apo-CaM within our system, as we measure bound versus free CaM without determining its specific orientation. Data from HEK293 cells indicate that approximately half of the CaM is Ca^2+^ free under resting conditions (Black *et al*., 2004), with the Ca^2+^-saturated CaM typically bound to other proteins (Wu & Bers, 2007). It is estimated that the concentration of free Ca^2+^-saturated CaM at similar resting conditions is approximately 50-60nM (Persechini & Cronk, 1999). Moreover, FGF12A has a reported K_d_ value of 107nM for CaM at its CaM-binding region (Mahling *et al*., 2021). Consequently, it is plausible that variations in the concentration of endogenous free Ca^2+^-saturated CaM at resting conditions could facilitate an interaction between FGF12A and endogenous CaM, which might not be detected by FRET imaging in HEK293 cells. This interaction could account for the effect of FGF12A on Nav1.5 at resting conditions, as depicted in Figure 6. In conclusion, these observations imply that FGF12A is responsive to alterations in [Ca^2+^]_i_ levels and the concentration of free CaM (both Ca^2+^-saturated and apo forms). Such changes could, in turn, modify the impact of FGF12A on the Nav1.5 channel.

As previously discussed, WT FGF12A is sensitive to the Ca^2+^ binding state of CaM, which influences the structural configuration of the Nav1.5:FGF12A complex and subsequently affects the inactivation of the Nav1.5 channel. In cardiomyocytes, the cardiac sodium channel Nav1.5 experiences cyclic changes in [Ca^2+^]_i_ levels. These fluctuations in [Ca^2+^]_i_ concentration lead to corresponding changes in free Ca^2+^-saturated CaM levels, increasing notably during cell contraction (Maier *et al*., 2006). With rising levels of free Ca^2+^-saturated CaM, Ca^2+^-CaM binds to the N-terminus of the FGF12A:Nav1.5 complex. This binding plays a crucial role in ensuring that the Nav1.5 channel inactivates correctly at the typical membrane potential, maintaining the integrity of the cardiac action potential. In neurons, the intracellular calcium transient is significantly shorter than in cardiomyocytes, lasting only a few milliseconds following neuronal action potentials (Llinás *et al*., 1981). Despite the brief Ca^2+^ flux, CaM remains activated (Ca^2+^-saturated) for a longer period, enabling it to signal other downstream effects well after the initial Ca^2+^ surge (Milikan *et al*., 2002). Consequently, Ca^2+^-saturated CaM can similarly bind to FGF12A and influence voltage-gated sodium channels like Nav1.5, following increased Ca^2+^ fluxes. This raises important questions regarding the on/off rate of CaM and FGF12A interactions and whether these interactions are dynamic, varying based on the relative concentration of free CaM available. Understanding these dynamics will be essential for comprehending how changes in [Ca^2+^]_i_ and CaM states impact the function of sodium channels in both cardiac and neuronal cells.

### Study Limitations

Due to the nature of FRET imaging, manual patch-clamping, and the ionomycin treatment, we were unable to measure the precise intracellular calcium concentration during recordings. For the purposes of this study, we limited ourselves to three levels of calcium concentrations: low (chelated with BAPTA), control (resting/basal levels), and high (elevated with KR solution and ionomycin). Our aim was not to mimic exact physiological levels of calcium concentrations, but instead to ensure that if any calcium related effects can be applied to the sodium channel complex, we could detect it. This required examining the extreme ends of the spectrum within living cells. Further work is needed to recapitulate our findings in more native environments, specifically at looking at the potential changes in the sodium channel complex throughout the calcium transient associated with excitable cells, potentially through CRISPR edited iPSC-derived cells.

## Appendix

### Control experiments to evaluate stoichiometry via FRET

In Ben-Johny et al. (2016), the authors showed that increasing Ca²⁺ concentrations alters the stoichiometric ratio of CaM to the Cav1.2 calcium channel, but not to the Nav1.5 sodium channel (Ben-Johny *et al*., 2016). To extend this framework, we employed a custom FRET imaging system (**Figure A1**) designed to quantify donor-to-acceptor ratios (Kang *et al*., 2023a). This system collected three signals: S(Donor)- 440nm excitation and 475nm emission, S(FRET)- 440nm excitation and 543nm emission, and S(Acceptor)- 510nm excitation and 543nm emission; from which we calculate the apparent donor- and acceptor-centric FRET efficiencies (ED and EA). Their relationship to the true FRET efficiency (E_FRET_) is described by Equations 1 and 2 where A_b_ and D_b_ represent the fractions of bound acceptors and donors, respectively (see Methods and Kang et al. 2023).

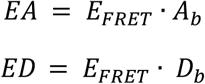

As both EA and ED scale with the proportion of bound donors and acceptors (donor fluorophores bound to acceptor fluorophores and vice versa), comparing them allows us to infer the ratio of acceptors to donors. When donors and acceptors are present in equal numbers, EA and ED converge, as shown by Ben-Johny et al. (2016) and Kang et al. (2023) (**Figure A1B**). Conversely, a shift in either EA or ED reflects changes in acceptor-to-donor ratios (**Figure A1C**).

To validate measuring the stoichiometry of auxiliary subunit via FRET imaging on our system we measured FRET pairs with the KCNQ1 channel, which is known to bind CaM at a 1:1 ratio46. Co-transfection of CFP-tagged KCNQ1 with YFP-tagged CaM at this ratio yielded equivalent EA and ED values, confirming the expected stoichiometry (**Figure A2A**). Adding a YFP tag to the extracellular region of KCNQ1 (S1-S2 linker) allowed for the presence of both an acceptor and donor on the KCNQ1 channel. In this configuration, CaM-CFP produced a twofold increase in EA relative to ED, indicative of a 1:2 acceptor:donor ratio (**Figure A2B**). Inversely, when co-transfected with CaM-YFP, the ratio was reversed and ED doubled relative to EA (**Figure A2C**). Together, these experiments demonstrate that our FRET imaging system can reliably detect ion channel and auxiliary subunit stoichiometry.

## Additional Information Figures

**Figure A1:**
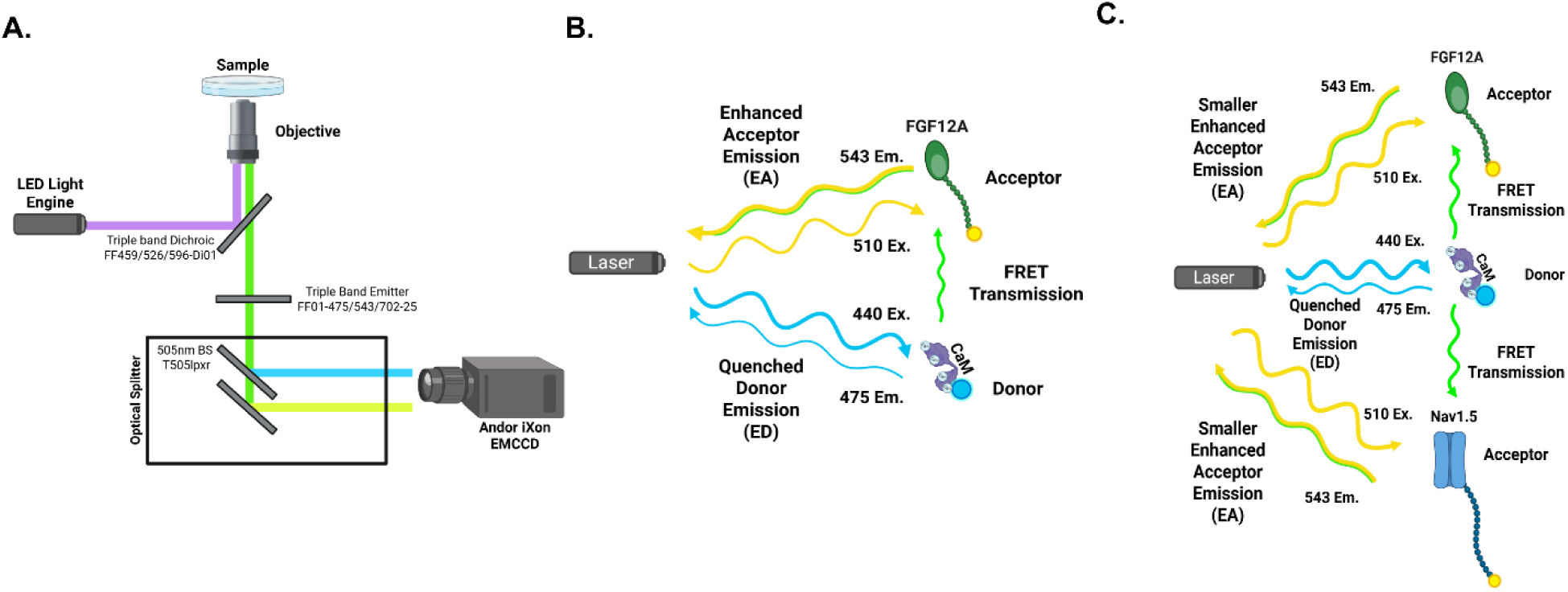
Donor (CaM-CFP) and acceptor (FGF12A-YFP, Nav1.5-YFP) ratios were assessed by comparing donor-(ED) and acceptor-centric (EA) FRET efficiencies. **(A)** Microscope setup for simultaneous multi-wavelength detection. **(B)** A 1:1 **EA-ED** relationship indicates equal donor and acceptor levels. **(C)** Higher **ED** than EA reflects excess acceptors **in** the complex.

**Figure A2:**
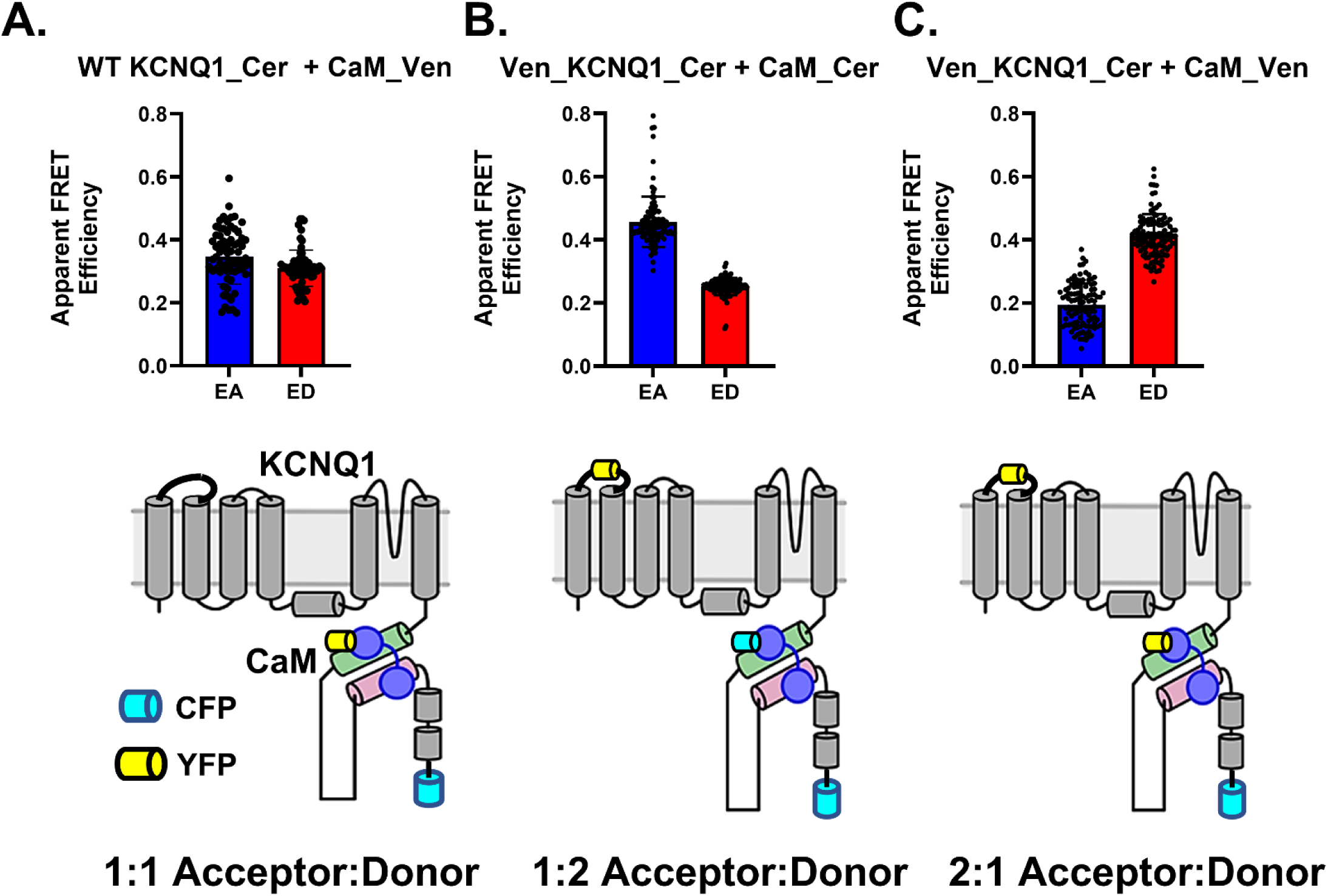
Live-cell FRET imaging validates measurement of donor-to-acceptor ratios. **(A)** KCNQ1-CaM complexes display a 1:1 ratio: YFP-CaM (acceptor) with CFP-KCNQ1 (donor) yields equivalent donor- and acceptor-centric FRET efficiencies (ED and EA). **(B)** Introducing an additional YFP tag on the KCNQ1 S1-S2 loop creates a dual-labeled KCNQ1 (CFP and YFP). Pairing this with CFP-CaM produces a 1:2 acceptor:donor ratio, reflected by higher EA relative to ED. (C) Adding YFP-CaM shifts the balance to a 2:1 acceptor:donor ratio.

**Figure A3:**
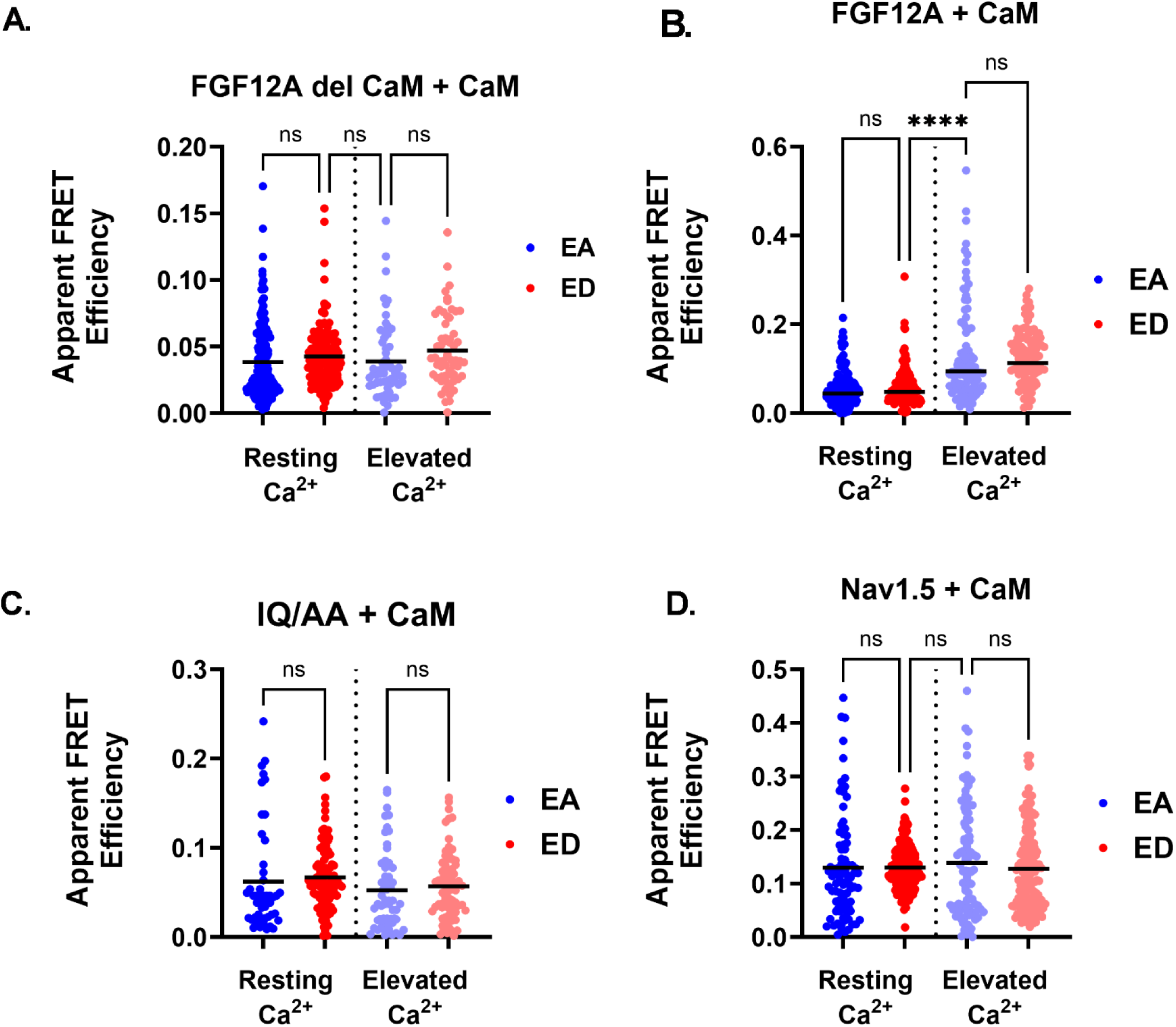
Scatter plots with mean values of apparent FRET efficiencies. Across all conditions: **(A)** FGF12A del CaM + **CaM,(B)** FGF12A + CaM, **(C)** IQ/AA+ CaM, and **(D)** Nav1.5 + CaM, no significant differences were observed between donor (ED) and acceptor-centric (EA) values, consistent with a 1:1 donor-to-acceptor ratio. The only condition showing increased FRET efficiency with elevated Ca^2^+ was FGF12A + CaM (****p < 0.0001). Some datasets overlap with those in Figure 2.

**Figure A4:**
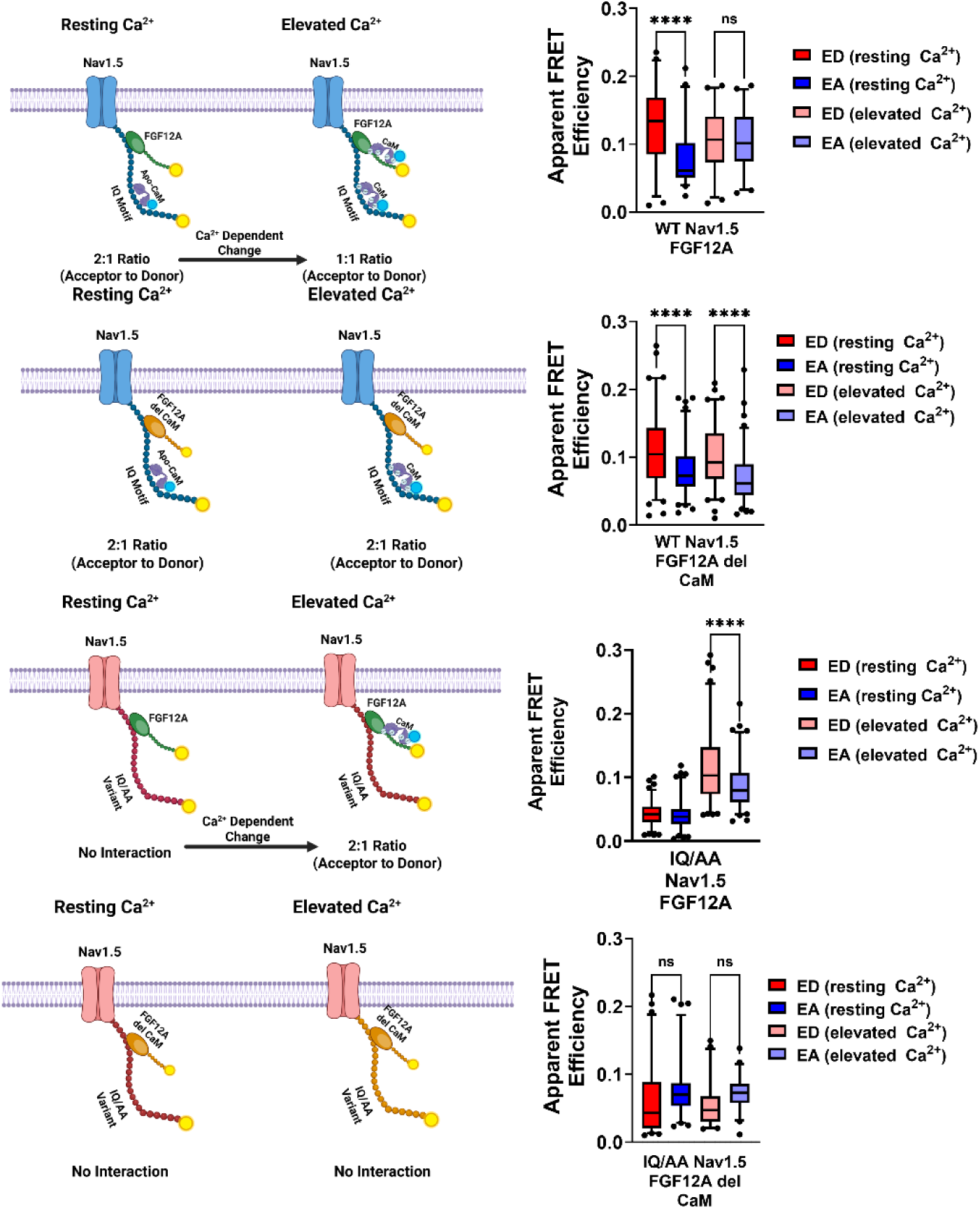
FRET analysis of Nav1.5:CaM:FGF12A interactions under varying calcium conditions. **(A)** At resting [Ca^2+^]i, WT Navi .5 shows an EA:ED ratio of 1:2, consistent with a single apo-CaM bound to the Nav1.5 CTD, as FGFI2A:CaM interactions require Ca^2+^. At elevated Ca^2+^, EA equals ED, reflecting a 1:1 ratio with two CaM molecules bound simultaneously to the Navi 5 CTD and FGF12A (“**p < 0.0001). (B) Deletion of the CaM-binding site in FGF12A abolished Ca^2+^-dependeπt binding (****p < 0.0001). (C) With the Navi .5 IQ/AA variant, CaM cannot bind the CTD, leaving only Ca^2+^-bound CaM to interact with FGF12A (“p = 0.012). (D) No significant FRET signal was detected when all CaM-binding sites are removed.

## Acknowledgements

This work was completed in its entirety within the Silva lab at Washington University in St. Louis. It was funded by NIH-NHLBI RO1 HL136553, NIH-NHLBI RO1HL150637-01A1 and the individual predoctoral fellowship 1F31HL164080-01A1.

